# Energy efficient synaptic plasticity

**DOI:** 10.1101/714055

**Authors:** Ho Ling Li, Mark C. W. van Rossum

## Abstract

Many aspects of the brain’s design can be understood as the result of evolutionary drive towards efficient use of metabolic energy. In addition to the energetic costs of neural computation and transmission, experimental evidence indicates that synaptic plasticity is metabolically demanding as well. As synaptic plasticity is crucial for learning, we examine how these metabolic costs enter in learning. We find that when synaptic plasticity rules are naively implemented, training neural networks requires extremely large amounts of energy when storing many patterns. We propose that this is avoided by precisely balancing labile forms of synaptic plasticity with more stable forms. This algorithm, termed synaptic caching, boosts energy efficiency manifold. Our results yield a novel interpretation of the multiple forms of neural synaptic plasticity observed experimentally, including synaptic tagging and capture phenomena. Furthermore our results are relevant for energy efficient neuromorphic designs.

---

The human brain only weighs 2% of the total body mass, but is responsible for 20% of resting metabolism [1, 2]. The brain’s energy need is believed to have shaped many aspects of its design, such as its sparse coding strategy [3, 4], the biophysics of the mammalian action potential [5, 6], and synaptic failure [7, 2]. As the connections in the brain are adaptive, one can design synaptic plasticity rules that further reduce the energy required for information transmission, for instance by sparsifying connectivity [8]. But in addition to the costs associated to neural information processing, experimental evidence suggests that memory formation, presumably corresponding to synaptic plasticity, is itself an energetically expensive process as well [9, 10, 11, 12].

To estimate the amount of energy required for plasticity, Mery and Kawecki [9] subjected fruit flies to associative conditioning spaced out in time, resulting in long-term memory formation. After training, the fly’s food supply was cut off. Flies exposed to the conditioning died some 20% quicker than control flies. Similarly, fruit flies doubled their sucrose consumption during the formation of aversive long-term memory [12], while forcing starving fruit flies to form such memories reduced lifespan by 30% [10]. Notably, less permanent forms of learning that don’t require protein synthesis have been observed to be energetically less costly [9, 10].

Motivated by these experimental results, we analyze the metabolic energy required to form associative memories in neuronal networks. We demonstrate that traditional learning algorithms are metabolically highly inefficient. Therefore we introduce a synaptic caching algorithm that is consistent with synaptic consolidation experiments, and distributes learning over transient and persistent synaptic changes. This algorithm increases efficiency manifold. Synaptic caching yields a novel interpretation to various aspects of synaptic physiology, and suggests more energy efficient neuromorphic designs.

## Results

To examine the metabolic energy cost associated to synaptic plasticity, we first study the perceptron. A perceptron is a single artificial neuron that attempts to binary classify input patterns. It forms the core of many artificial networks and has been used to model plasticity in cerebellar Purkinje cells. We consider the common case where the input patterns are random patterns each associated to a randomly chosen binary output. Upon presentation of a pattern, the perceptron output is calculated and compared to the desired output. The synaptic weights are modified according to the perceptron learning rule, Fig. 1A. This is repeated until all patterns are classified correctly [13, see Methods]. Typically, the learning takes multiple iterations over the whole dataset (‘epochs’).

**Figure 1:**
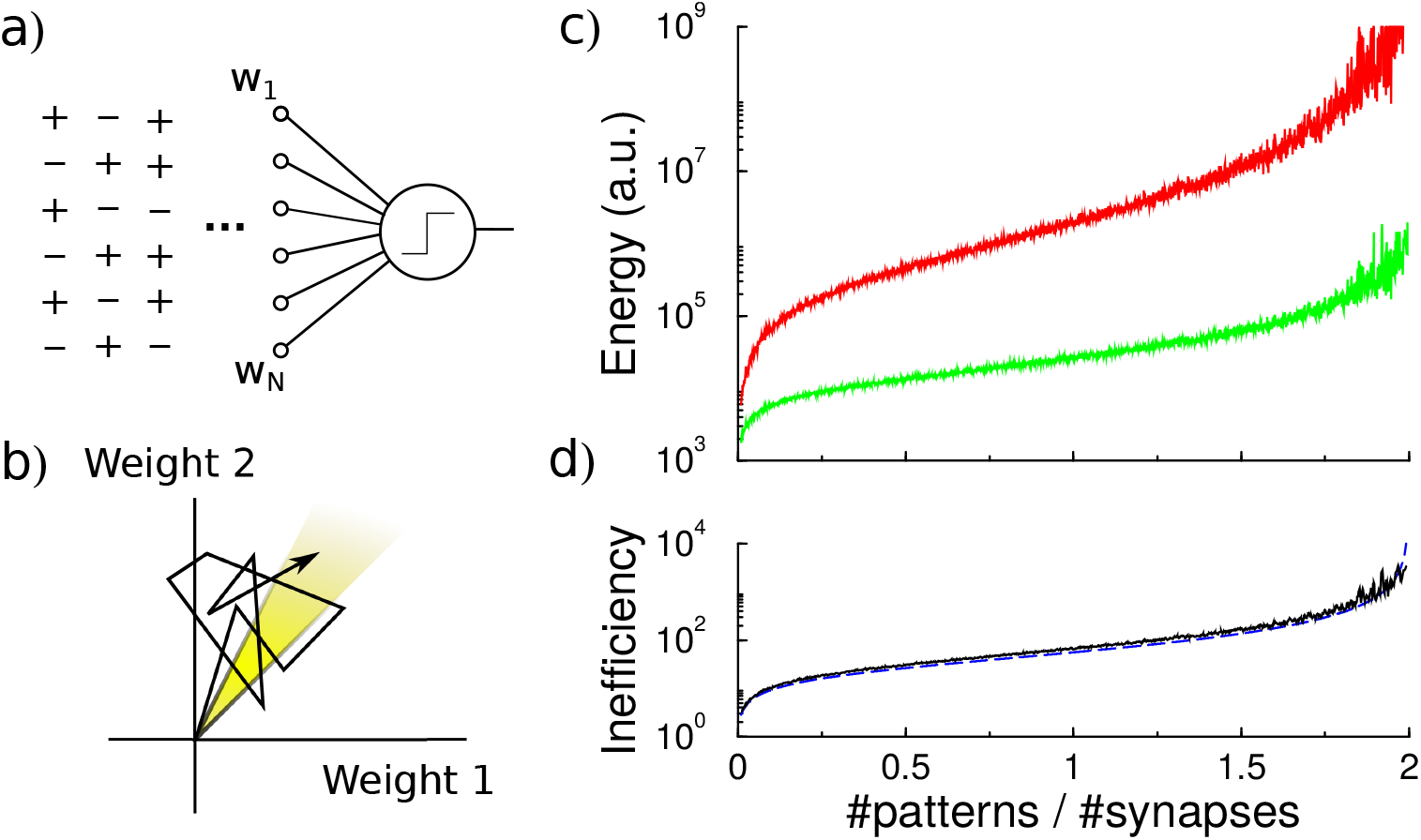
Energy efficiency of perceptron learning. (a) A perceptron cycles through the patterns and updates its synaptic weights until all patterns produce their correct target output. (b) During learning the synaptic weights follow approximately a random walk until they find the solution (yellow region). The energy consumed by the learning corresponds to the total length of the path (under the *L*_1_ norm). (c) The energy required to train the perceptron diverges when storing many patterns (red curve). The minimal energy required to reach the correct weight configuration is shown for comparison (green curve). (d) The inefficiency, defined as the ratio between actual and minimal energy plotted in panel c, diverges as well (black curve). The overlapping blue curve corresponds to Eq. 3 in the text.

As it is not well known how much metabolic energy is required to modify a biological synapse, and how this depends on the amount of change and the sign of the change, we propose a parsimonious model. We assume that the metabolic energy for every modification of a synaptic weight is proportional to the amount of change, no matter if this is positive or negative, although there is evidence that synaptic depression involves different pathways than synaptic potentiation, see e.g. [14]. The total metabolic cost *M* (in arbitrary units) to train a perceptron is

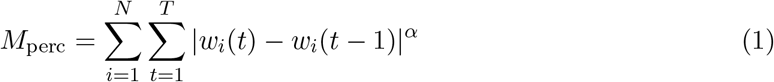

where *N* is the number of synapses, *w*_*i*_ denotes the synaptic weight at synapse *i*, and *T* is the total number of time-steps required to learn the classification. The exponent *α* is set to one, but our results below are similar whenever 0 ≤ *α* ≲ 2.

Learning can be understood as a search in the space of synaptic weights for a weight vector that leads to correct classification of all patterns, Fig. 1B. The synaptic weights approximately follow a random walk (Methods), and the energy is proportional to the length of this walk under the *L*_1_ norm, Eq. 1. The perceptron learning rule is energy inefficient, because repeatedly, weight modifications made to correctly classify one pattern are partly undone when learning another pattern, but as both processes require energy this is inefficient.

The energy required by the perceptron learning rule depends on the number of patterns *P* to be classified. The set of correct weights spans a cone in *N*-dimensional space (yellow region in Fig. 1B). As the number of patterns to be classified increases, the cone containing correct weights shrinks and the random walk becomes longer [15]. Near the critical capacity of the perceptron (*P* = 2*N*), the number of epochs required diverges as (2 − *P*/*N*)^−2^, [16]. The energy required, which is proportional to the number of updates that the weights undergo, follows a similar behavior, Fig. 1C.

It is useful to consider the theoretical minimal energy required to classify all patterns. The most energy efficient algorithm would somehow directly set the synaptic weights to their desired final values. Geometrically, the random walk trajectory of the synaptic weights to the target is replaced by a path straight to the correct weights. Given the initial weights *w*_*i*_(0) and the final weights *w*_*i*_(*T*), the energy required in this idealized case to set the synapses correctly is

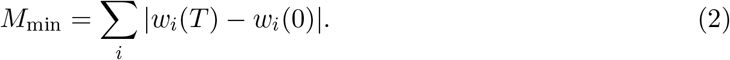

While the minimal energy also grows with the memory load (Methods), it increases less steeply, Fig. 1C.

We express the metabolic efficiency of a learning algorithm as the ratio between the energy the algorithm requires and the minimal energy (the gap between the two curves in Fig. 1C). As the number of patterns increases, the inefficiency of the perceptron rule rapidly grows, Fig. 1D, as (see Methods)

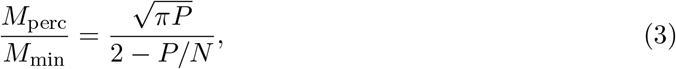

which fits the simulations well.

There is evidence that both cerebellar and cortical neurons are operating close to their maximal memory capacity [17, 18]. Indeed, it would appear wasteful if this were not the case. However, the above result demonstrates that for instance classifying 1900 patterns by a neuron with 1000 synapses with the traditional perceptron learning requires about ∼900 times more energy than minimally required. As the fruit-fly experiments indicate that even storing a single association in long-term memory is already metabolically expensive, storing many memories would thus require very large amounts of energy if the biology would naively implement these learning rules.

### Synaptic caching

How can the conflicting demands of energy efficiency and high storage capacity be met? The minimal energy argument presented above suggests a way to increase energy efficiency. There are forms of plasticity - anaesthesia resistant memory in flies and early-LTP/LTD in mammals - that decay and do not require protein synthesis. Such transient synaptic changes can be induced using a massed, instead of a spaced, stimulus presentation protocol. Fruit-fly experiments show that this form of plasticity is much less energy-demanding than long-term memory [9, 10, 12]. In mammals there is evidence that synaptic consolidation, but not transient plasticity, is suppressed under low energy conditions [19]. Inspired by these findings we propose that the transient form of plasticity constitutes a synaptic variable that accumulates the synaptic changes across multiple updates in a less expensive form of memory; only occasionally the changes are consolidated. We call this *synaptic caching*.

Specifically, we assume that each synapse is comprised of a transient component *s*_*i*_ and a persistent component *l*_*i*_. The total synaptic weight is their sum, *w*_*i*_ = *s*_*i*_ + *l*_*i*_. We implement synaptic caching as follows, Fig. 2A: For every presented pattern, changes in the synaptic strength are calculated according to the perceptron rule and are accumulated in the transient component that decays exponentially to zero. If, however, the absolute value of the transient component of a synapse exceeds a certain consolidation threshold, all synapses of that neuron are consolidated (vertical dashed line in Fig. 2A), the value of the transient component is added to the persistent weight, and the transient weight is reset to zero.

**Figure 2:**
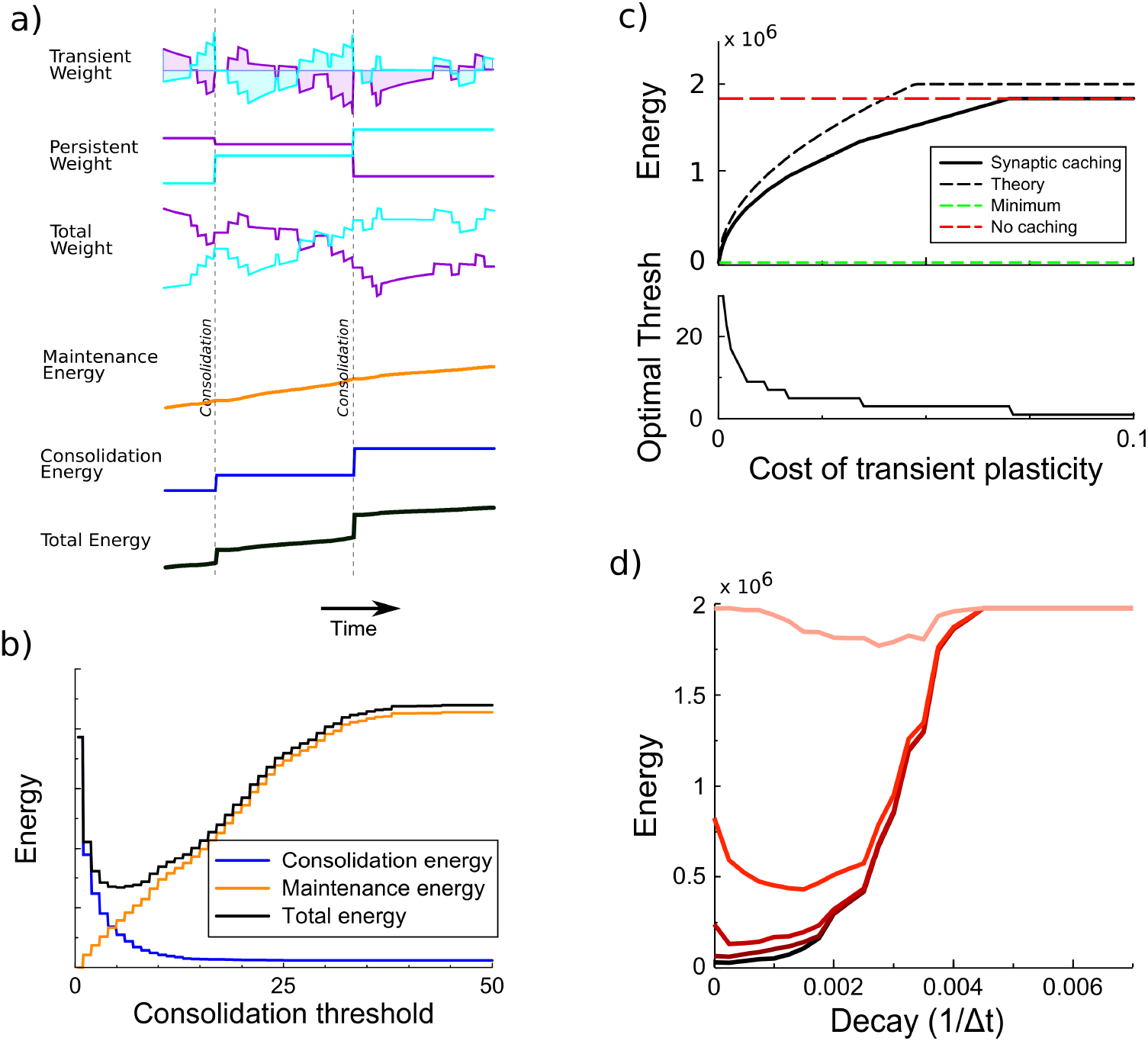
Synaptic caching algorithm. (a) Changes in the synaptic weights are initially stored in metabolically cheaper transient decaying weights. Here two example weight traces are shown (blue and magenta). The total synaptic weight is composed of transient and persistent forms. Whenever any of the transient weights exceed the consolidation threshold, the weights become persistent and the transient values are reset (vertical dashed line). The corresponding energy consumed during the learning process consists of two terms: the energy cost of maintenance is assumed to be equal to the magnitude of the transient weight (shaded area in top traces); energy cost for consolidation is incurred at consolidation events. (b) The total energy is composed of the energy to occasionally consolidate and the energy to support transient plasticity. Here it is minimal for an intermediate consolidation threshold. (c) The amount of energy required for learning with synaptic caching, in the absence of decay of the transient weights (black curve). When there is no decay and no maintenance cost the energy equals the minimal one (green line) and the efficiency gain is maximal. As the maintenance cost increases, the optimal consolidation threshold decreases (lower panel) and the total energy required increases, until no efficiency is gained at all by synaptic caching. (d) The amount of energy required for learning as a function of the decay of transient plasticity for various values of the maintenance cost (from bottom to top maintenance cost *c* = 0, 10^−4^, 10^−3^, 10^−2^, 10^−1^). Broadly, stronger decay will increase the energy required and hence reduce efficiency.

How much efficiency can be improved with synaptic caching depends on the limitations of transient plasticity. If the transient synaptic component could store information indefinitely at no metabolic cost, consolidation could be postponed until the end of learning and the energy would equal the minimal energy Eq. 2. Hence the efficiency gain would be maximal. However, we assume that the efficiency gain of synaptic caching is limited because of two effects: 1) The transient component decays exponentially (with a time-constant *τ*). 2) There might be a maintenance cost associated to maintaining the transient component. Biophysically, transient plasticity might correspond to an increased/decreased vesicle release rate [20, 21] so that it diverges from its optimal value [7].

To estimate the energy saved by synaptic caching we assume that the maintenance cost is proportional to the transient weight itself and incurred every time-step Δ*t* (shaded area in the top traces of Fig. 2A)

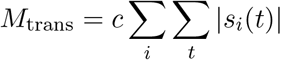

While experiments indicate that transient plasticity is metabolically far less demanding than the persistent form, the precise value of the maintenance cost is not known. We encode it in the constant *c*; the theory also includes the case that *c* is zero.

Next we need to include the energetic cost of consolidation. Currently it is unknown how different components of synaptic consolidation, such as signaling, protein synthesis, transport to the synapses and changing the synapse, contribute to this cost. We assume the metabolic cost to consolidate the synaptic weights is *M*_cons_ = Σ_*i*_Σ_*t*_ |*l*_*i*_(*t*) − *l*_*i*_(*t* − 1)|. The form of the consolidation energy is identical to Eq. 1, but in contrast to the standard perceptron learning, where synapses are consolidated every time a weight is updated, now changes in the persistent component *l*_*i*_ only occur when consolidation occurs. One can add a term similar to a one-off cost for changing the transient component, but as that would not vary with consolidation rate it is not included.

The energy gain achieved by synaptic caching depends on the consolidation threshold, Fig. 2B. When the threshold is low, consolidation occurs often and the energy approaches the one without synaptic caching. When on the other hand the consolidation threshold is high, the expensive consolidation process occurs rarely, but the maintenance cost of transient plasticity is high, moreover the decay will lead to forgetting of unconsolidated memories, slowing down learning and increasing the energy cost. Thus the consolidation energy decreases for larger thresholds, whereas the maintenance energy increases, Fig. 2B (see Methods). As a result of this trade-off there is an optimal threshold, which depends on the decay and the maintenance cost, that balances persistent and transient forms of plasticity. To analyze the efficiency gain we use this optimal value.

Fig. 2C shows the energy required to train the perceptron for the case when the transient component does not decay. When the maintenance cost is absent (*c* = 0), consolidation is best postponed until the end of the learning and the energy is as low as the theoretical minimal bound. As *c* increases, it becomes beneficial to consolidate more often, i.e. the optimal threshold decreases, Fig. 2C bottom panel. The required energy increases until the maintenance cost becomes so high that it is better to consolidate after every update and no energy is saved with synaptic caching. The efficiency is well described by analysis, Fig. 2C (Methods).

Fig. 2D examines the amount of savings as a function of the strength of the decay (expressed as 1/*τ*) of the transient component for various levels of maintenance cost. Efficiency is high when there is no decay. However, if the transient component decays it is best to consolidate more frequently, even when the maintenance cost is zero, as otherwise, information is lost and learning time increases. Interestingly, with intermediate amounts of decay somewhat less energy is required than without any decay. The reason is a slight reduction on number of epochs required when the synaptic weights decay.

In the above implementation of synaptic caching, consolidation of all synapses was triggered when transient plasticity at a single synapse exceeded a certain threshold. This resembles the synaptic tagging and capture phenomenon where plasticity induction leads to transient changes and sets a tag; only strong enough stimulation results in proteins being synthesized and being delivered to all tagged synapses, consolidating the changes [22, 23]. There are a number of alternative ways to model the interaction between synapses: the threshold could be synapse-specific or neuron-wide, and the consolidation could be synapse-specific or neuron-wide, Fig. 3A. In practice there are three possibilities: First, consolidation might be set to occur whenever transient plasticity at a synapse crosses the threshold and only that synapse is consolidated. Second, a hypothetical signal might send to the soma and consolidation of all synapses occurs once transient plasticity at any synapse crosses the threshold (used in Figs. 2 and 4). Thirdly, a hypothetical signal might be accumulated in or near the soma and consolidation of all synapses occurs once this total transient plasticity across synapses crosses the threshold. Only cases 2 and 3 are consistent with synaptic tagging and capture experiments, where consolidation of one synapse also leads to consolidation of another synapse that would otherwise decay back to baseline [22, 24]. Notably, all variants lead to comparable efficiency gains, Fig. 3B.

**Figure 3:**
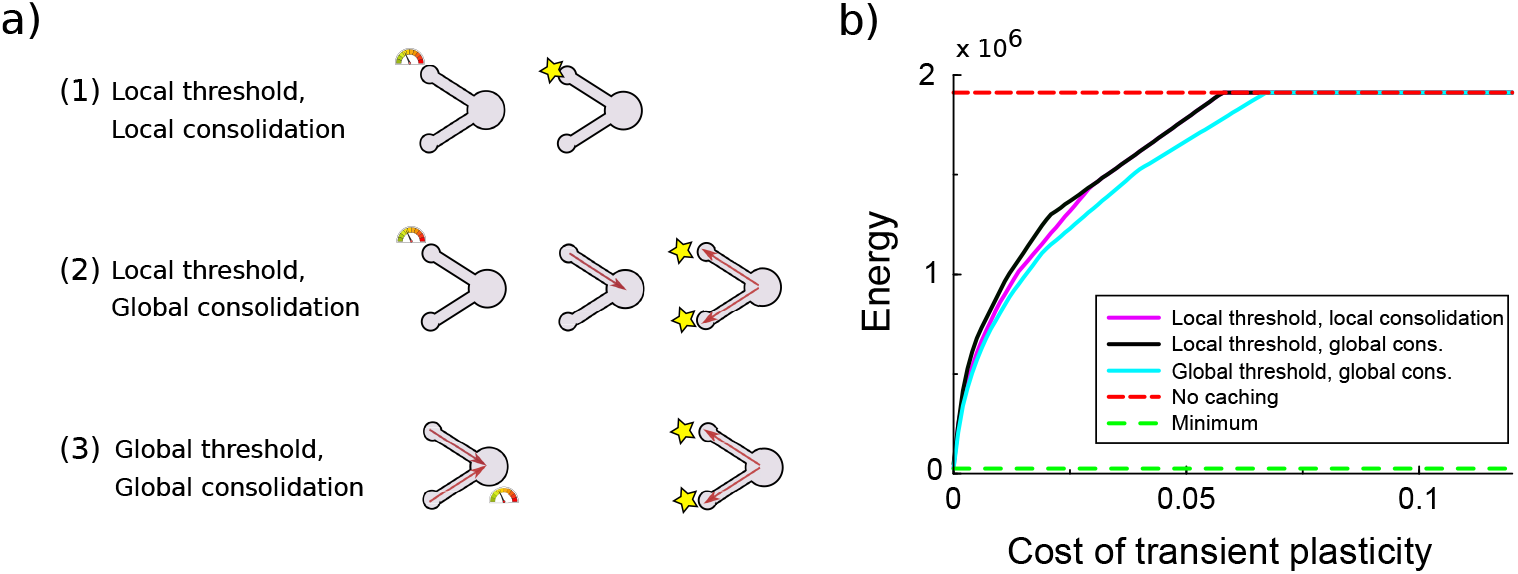
Comparison of various variants of the synaptic caching algorithm. (a) Schematic representation of variants to decide when consolidation occurs. From top to bottom: 1) Consolidation (indicated by the star) occurs whenever transient plasticity at a synapse crosses the consolidation threshold and only that synapse is consolidated. 2) Consolidation of all synapses occurs once transient plasticity at any synapse crosses the threshold. 3) Consolidation of all synapses occurs once the total transient plasticity across synapses crosses the threshold. (b) Energy required to teach the perceptron is comparable across algorithm variants. Consolidation thresholds were optimized for each algorithm and each maintenance cost of transient plasticity individually. In this simulation the transient plasticity did not decay.

**Figure 4:**
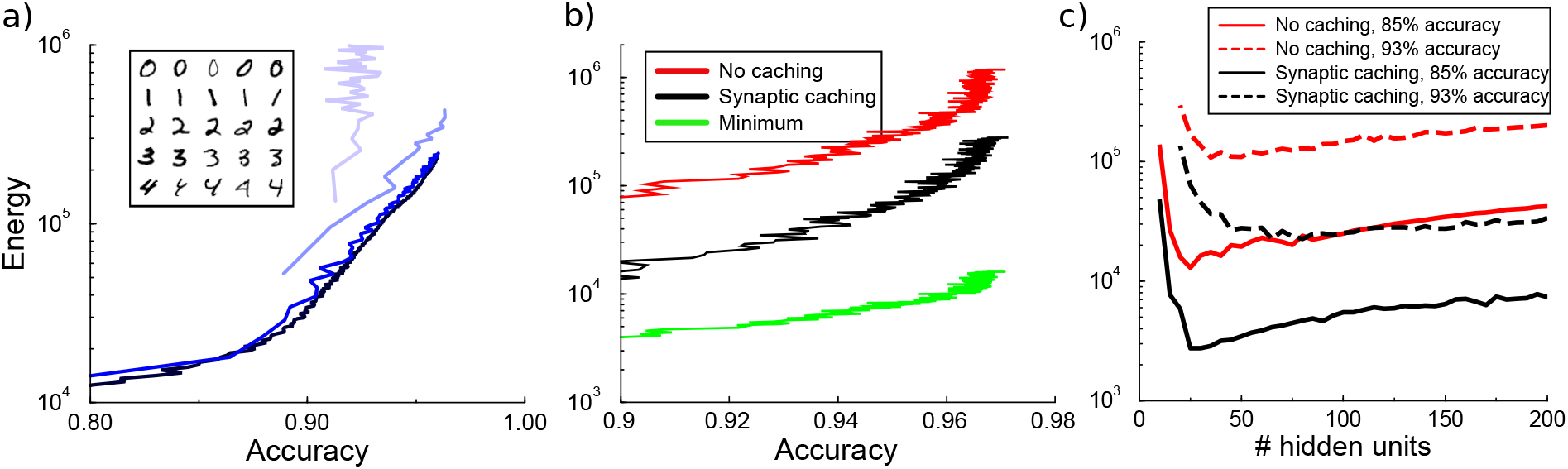
Energy cost to train a multi-layer back-propagation network to classify digits from the MNIST data set. (a) Energy rises with the accuracy of identifying the digits from a held-out test data. Except for the larger learning rates, the energy is independent of the learning rate (from bottom to top learning rate *η* = 10^−3^, 10^−2^, 10^−1^, 0.5). Inset shows some MNIST examples. (b) Comparison of energy required to train the network with/without synaptic caching, and the minimal energy. As for the perceptron and depending on the cost of transient plasticity, synaptic caching can reduce energy need manifold. (c) The impact of number of hidden units in the network with back-propagation on the metabolic cost. The network is trained to classify digits from the MNIST dataset to 85% and 93% accuracy. Both with and without synaptic caching, energy needs are high when the number of hidden units is barely sufficient. Parameters for transient plasticy in (b) and (c): *τ* = 1000, *c* = 0.001.

In summary we see that synaptic caching can in principle achieve large efficiency gains, bringing efficiency close to the theoretical minimum.

### Energy of learning in multi-layer network

Since the perceptron is a rather restrictive framework, we wondered whether the efficiency gain of synaptic caching can be transferred to multi-layer networks. Therefore we implement a multi-layer network trained with back-propagation. Back-propagation networks learn the associations of patterns by approaching the minimum of the error function through stochastic gradient descent. We use a network with one hidden layer with by default 100 units to classify hand-written digits from the MNIST dataset. As we train the network, we intermittently interrupt the learning to measure the energy consumed for plasticity and measure the performance on a held-out test-set. This yields a curve relating energy to accuracy.

Similar to a perceptron, learning without synaptic caching is metabolically expensive in a back-propagation network. Until reaching maximal accuracy, energy rises approximately exponentially with accuracy, after which additional energy do not lead to further improvement. When the learning rate is sufficiently small, the metabolic cost of plasticity is independent of the learning rate. At larger learning rates, learning no longer converges and energy goes up steeply without an increase in accuracy, Fig. 4A. With the exception of these large rates, these results show that changing the learning rate does not save energy.

Similar to the perceptron, we evaluate how much energy would be required to directly set the synaptic weights to their final values. Traditional learning without synaptic caching is once again energetically inefficient, expending at least ∼ 20 times more energy compared to this theoretical minimum whatever the desired accuracy level is, Fig. 4B. However, by splitting the weights into persistent synaptic weights and transient synaptic caching weights, the network can save substantial amounts of energy. As for the perceptron, depending on the decay and the maintenance cost the energy ranges from as little as the minimum to as much as the energy required without caching. Thus the efficiency gain of synaptic caching found for the perceptron carries over to multi-layer networks.

It might seem that smaller networks would be metabolically less costly, because small networks simply contain fewer synapses to modify. On the other hand, for the perceptron metabolic costs rise rapidly when cramming many patterns into it. We wondered therefore how energy cost depends on network size in the multi-layer network. Since the number of input units is fixed to the image size and the number of output units equals the ten output categories, we adjust the number of hidden units. As expected, higher accuracies require more hidden units and energy, Fig. 4C. The network fails to reach the desired accuracy if the number of hidden units is too small. When the network size is barely above the minimum requirement, the network has to compensate the lack of hidden units with longer training time and hence a larger energy expenditure. However, very large networks also require more energy. These results show that from an energy perspective there exists an optimal number of neurons to participate in memory formation.

## Discussion

Experiments on formation of a long-term memory of a single association suggest that synaptic plasticity is an energetically expensive process. We have shown that energy requirements rise steeply as memory load or designated accuracy level increase. This indicates trade-offs between energy consumption, and network capacity and performance. To improve efficiency we have proposed an algorithm named synaptic caching: temporarily storing changes in the synaptic strength at the transient forms of plasticity, which are, determined by a threshold, only occasionally consolidated to the persistent forms. Depending on the characteristics (decay and maintenance cost) of transient plasticity, this can lead to large energy savings in the energy required for synaptic plasticity. Further savings might be possible by adjusting the consolidation threshold as learning progresses and by being pathway-specific [25].

The implementation of a consolidation threshold is similar to what has been observed in physiology, in particular in the synaptic tagging and capture literature [26]. Our results thus give a novel interpretation of those findings. Synaptic consolidation is known to be affected by reward, novelty and punishment [26], which is compatible with a metabolic perspective as energy is expended only when the stimulus is worth remembering. In addition, our results for instance explain why consolidation is competitive, but transient plasticity is less so [27], namely the formation of long-term memory is precious. Consistent with this, there is evidence that encouraging consolidation increases energy consumption [12]. We also predict that the transient weight changes act as an accumulative threshold for consolidation. That is, sufficient transient plasticity should trigger consolidation, even in the absence of other consolidation triggers. Future characterization of the energy budget of synaptic plasticity should allow more precise predictions of our theory.

Combining persistent and transient storage mechanisms is a strategy well known in traditional computer systems to provide a faster and often energetically cheaper access to memory. In computer systems permanent storage of memories typically requires transmission of all information across multiple transient cache systems until reaching a long-term storage device and the transfer of information can often be a bottleneck in computer architectures and consumes considerable power in modern computers [28]. However, in the nervous system transient and persistent synapses appear to exist next to each other. The consolidation of information in a synapse does not require moving that information. Using this setup, biology appears to have found a more efficient way to store information.

Memory stability has long fascinated researchers [29], and in some cases forgetting can be beneficial [30]. Here we argue that the main benefit of more transient forms of plasticity is to permit the network to explore the weight space to find a desirable weight configuration using less energy. Besides suggesting forms of plasticity with different persistence, the cost of synaptic plasticity could potentially have influenced other aspects of neurobiological design. In principle, homeostasis and long-term stability could impact the cost of learning as well. Moreover, this work focuses on just the metabolic cost of synaptic plasticity, but the brain also expends significant amounts of energy on spiking, synaptic transmission, and maintaining resting potential. Further study is needed to understand how this impacts total energy cost during and after learning.

## Methods

### Energy efficiency of the perceptron

For perceptron we can calculate the energy efficiency of both the classical perceptron and the gain achieved by synaptic caching. We first consider the case that transient plasticity does not decay, as this allows important theoretical simplifications. In the perceptron learning to classify binary patterns Eq. 7, the weight updates are either +*η* or −*η*, where *η* is the learning rate, so that the energy spent Eq. 1 per update per synapse equals *η*. Hence the total energy spent to classify all patterns *M*_perc_ = *NKη*, where *K* is the total number of updates. We find numerically that *K* = 2*P*/(2 − *P*/*N*)^2^.

To calculate the efficiency we compare this to the minimal energy necessary to reach the final weight vector in the perceptron. We approximate the weight trajectory followed by the perceptron algorithm by a random walk. After *K* updates of step-size *η* the weights approximate a Gaussian distribution with zero mean and variance *Kη*^2^. In simulations the variance in the weights is about 20% smaller, likely reflecting correlations in the learning process not captured in the random walk approximation. By short-cutting the random walk, the minimal energy required to reach the weight vector is 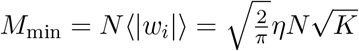. Hence, we find for the inefficiency (see Fig. 1D)

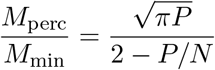

### Efficiency of synaptic caching

To calculate the efficiency gained with synaptic caching we need to calculate both the consolidation energy and the maintenance energy. The consolidation energy equals the number of consolidation events times the size of the updates. The size of the weight updates is equal to the consolidation threshold *θ*, while the number of consolidation events follows from a random walk argument as *NK*(⌈*θ*/*η*⌉)^2^. The ceiling function expresses the fact that when the threshold is smaller than learning rate, consolidation will always occur; we temporarily ignore this scenario. In addition, at the end of learning all remaining transient plasticity is consolidated, which requires an energy *N* 〈|*s*_*i*_(*T*)|〉. Assuming that the probability distribution *P*(*s*) has reached steady state, it has a triangular shape (see below) and 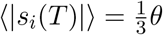 so that the total consolidation energy

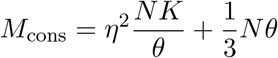

The transient energy is (again assuming that *P* (*s*) has reached steady state)

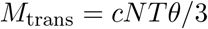

where *T* is the number of time-steps required for learning. Using that 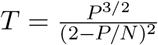, the total energy when using synaptic caching is 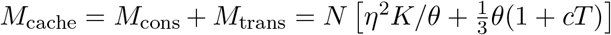. The optimal threshold 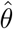 is given by 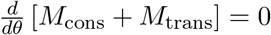 or

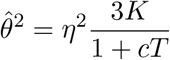

at which the energy is 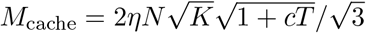. And so the efficiency of synaptic caching is 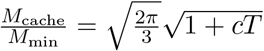. However, as consolidation can maximally occur only once per time-step, *M*_cons_ cannot exceed *M*_perc_ so that the inefficiency is

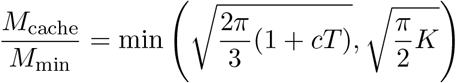

This equation reasonably matches the simulations, Fig. 2C (labeled ‘theory’).

### Decaying transient plasticity

When transient plasticity decays, the situation is more complicated as the learning time depends on the strength of the decay and to our knowledge no analytical expression exists for it. However, it is still possible to estimate the power, i.e. the energy per time unit, for both the transient component, denoted *m*_trans_, and the consolidation component, *m*_cons_. Under the random walk approximation every time the perceptron output does not match the desired output, the transient weight *s*_*i*_ is updated with an amount Δ*s*_*i*_ drawn from a distribution *Q*, with zero mean and variance *σ*^2^. Given the update probability *p*, i.e. the fraction of patterns not yet classified correctly, one has *Q*_*s*_(*η*) = *Q*_*s*_(−*η*) = *p*/2 and *Q*_*s*_(0) = 1 − *p*, so that 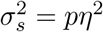. We assume that the number of updates slowly decreases as learning progresses, hence *p* is quasi-stationary.

Every time-step Δ*t* = 1 the transient weights decay with a time-constant *τ*. The synapse is consolidated and *s*_*i*_ is reset to zero whenever the absolute value of the caching weight |*s*_*i*_| exceeds *θ*. Given *p* and *τ*, we would like to know: 1) how often consolidation events occur which gives consolidation power and 2) the maintenance power *m*_trans_ = *cN* 〈|*s*_*i*_|〉. This problem is similar to the random walk to threshold model used for integrate-and-fire neurons, but here there are two thresholds: *θ* and −*θ*.

Under the assumptions of small updates and a smooth resulting distribution, the evolution of the probability distribution *P*(*s*_*i*_) is described by the Fokker-Planck equation, which in the steady state gives

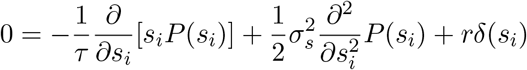

The last term is a source term that describes the re-insertion of weights by the reset process. The boundary conditions are *P* (*s*_*i*_ = ±*θ*) = 0. While *P*(*s*_*i*_) is continuous in *s*_*i*_, the source introduces a cusp in *P*(*s*_*i*_) at the reset value. Conservation of probability ensures that *r* equals the outgoing flux at the boundaries. One finds

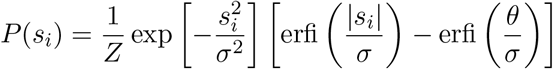

where erfi(*x*) = −*i*erf(*ix*), 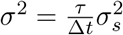 and with normalization factor

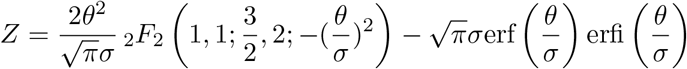

where _2_*F*_2_ is the generalized hypergeometric function. In the limit of no decay this becomes a triangular distribution *P*(*s*_*i*_) = [*θ* − |*s*_*i*_|]/*θ*^2^.

We obtain maintenance power

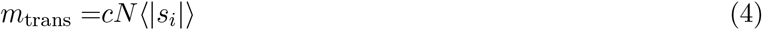

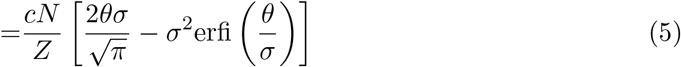

For small *θ*/*σ*, i.e. small decay, this is linear in *θ*, 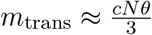. It saturates for large *θ* because then the decay dominates and the threshold is hardly ever reached.

The consolidation rate follows from Fick’s law

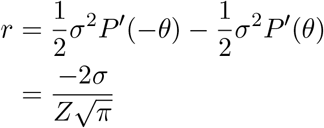

The consolidation power is

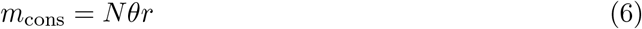

In the limit of no decay one has *r* = *σ*^2^/*θ*^2^, so that *m*_cons_ = *pNη*^2^/*θ*. Strictly speaking this approximates learning with a random walk process and assumes local consolidation, Fig. 3A. However, Eqs. 5 and 6 give a good prediction of the simulation when provided with the time-varying update probability from the simulation, Fig. 5.

**Figure 5:**
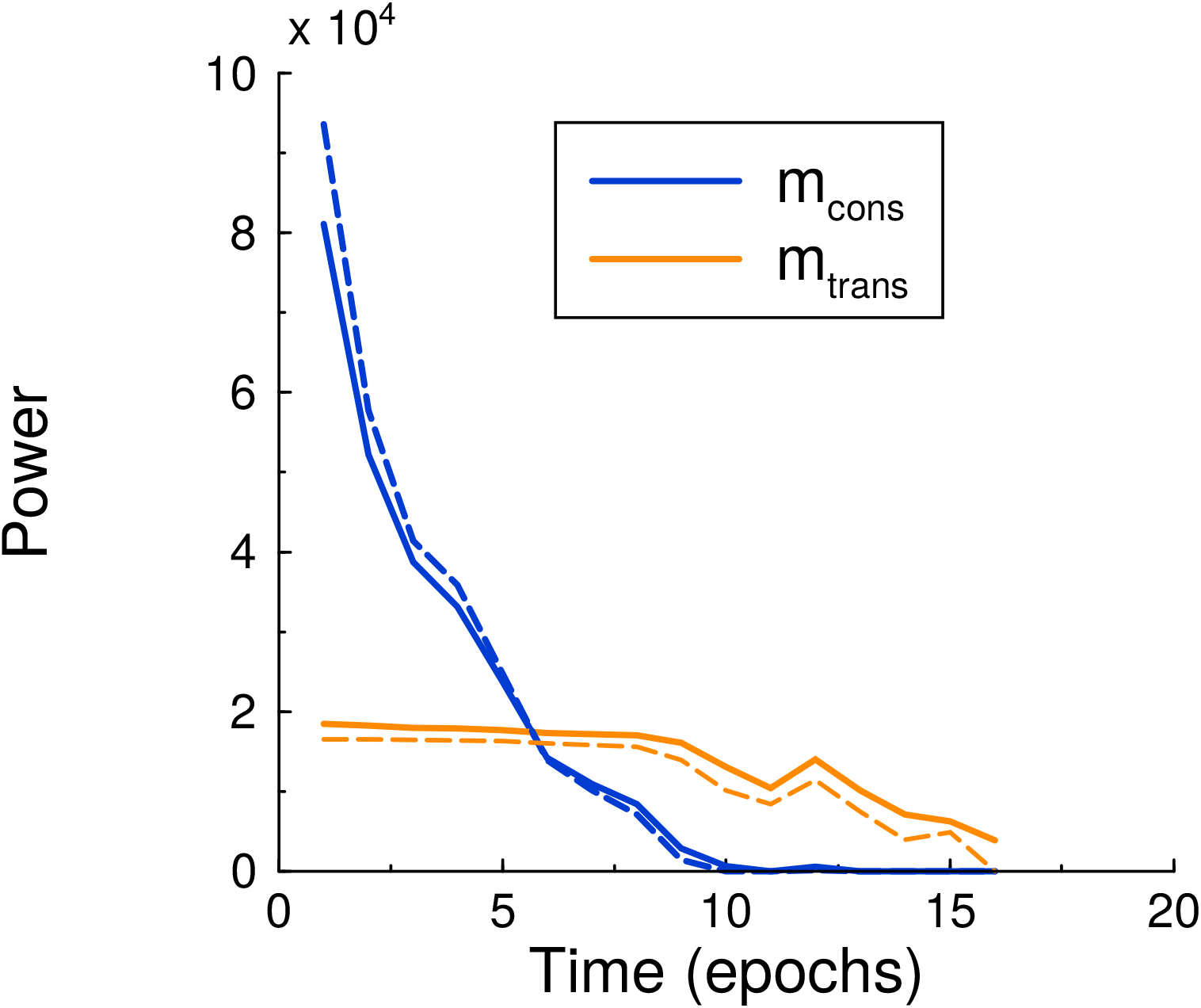
Maintenance and consolidation power. Power (energy per epoch) of the perceptron vs epoch. Solid curves are from simulation, dashed curves are the theoretical predictions, Eqs. 5 and 6, with their *σ* calculated by using the perceptron update rate *p* extracted from the simulation. Both powers are well described by the theory. Parameters: *τ* = 500, *c* = 0.01, *θ* = 5.

### Simulations

#### Perceptron

Unless stated otherwise, we use a perceptron with *N* = 1000 input units to classify *P* = *N* random binary (±1 with equal probability) input patterns ***x***^(*p*)^, each to be associated to a randomly assigned desired binary output *d*^(*p*)^. Each input unit is connected with a weight *w*_*i*_ signifying the strength of the connection. An ‘always-on’ bias unit with corresponding weight is included to adjust the threshold of the perceptron. The perceptron output *y* of a pattern is determined by the Heaviside step function Θ, *y* = Θ(w.*x*). If for a given pattern *p*, the output does not match the desired pattern output, ***w*** is adjusted according to

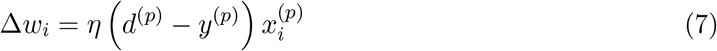

where the learning rate *η* can be set to one without loss of generality. The perceptron algorithm cycles through all patterns until classified correctly. In principle the magnitude of the weight vector, and hence the minimal energy, can be arbitrarily small for a noise-free binary perceptron. However, this paradox is resolved as soon as robustness to any post-synaptic noise is required.

#### Multi-layer networks

For the multi-layer networks trained on MNIST, we use networks with one hidden layer, logistic units, and one-hot encoding at the output. Weights are updated according to the mean squared error back-propagation rule without regularization.

## Acknowledgments

This project is supported by the Leverhulme Trust with grant number RPG-2017-404. MvR is supported by Engineering and Physical Sciences Research Council (EPSRC) grant EP/R030952/1. We would like to thank Joao Sacramento and Simon Laughlin for discussion and inputs.

## Author contribution

MvR designed the experiment. HLL and MvR did the simulations and calculations, and co-wrote the paper.

